# Structural basis for tuning activity and membrane specificity of bacterial cytolysins

**DOI:** 10.1101/2020.06.16.154724

**Authors:** Nita R. Shah, Tomas B. Voisin, Edward S. Parsons, Courtney M. Boyd, Bart W. Hoogenboom, Doryen Bubeck

## Abstract

Cholesterol-dependent cytolysins (CDCs) form protein nanopores to lyse cells. They target eukaryotic cells using different mechanisms, but all require the presence of cholesterol to pierce lipid bilayers. How CDCs use cholesterol to selectively lyse cells is essential for understanding virulence strategies of several pathogenic bacteria, and for repurposing CDCs to kill new cellular targets. Here we address that question by trapping an early state of pore formation for the CDC intermedilysin, bound to the human immune receptor CD59 in a nanodisc model membrane. Our cryo-electron microscopy map reveals structural transitions required for oligomerization, which include the lateral movement of a key amphipathic helix. We demonstrate that the charge of this helix is crucial for tuning lytic activity of CDCs. Furthermore, we discover modifications that overcome the requirement of cholesterol for membrane rupture, which will facilitate engineering the target-cell specificity of pore-forming proteins.

## INTRODUCTION

Pore-forming proteins rupture lipid bilayers to kill target cells. They comprise the largest class of virulence factors for pathogenic bacteria and are prevalent in all kingdoms of life^1^. Cholesterol-dependent cytolysins (CDCs) are pore-forming proteins secreted by more than five genera of Gram-positive bacteria, including human pathogens from *Streptococcus, Clostridium*, and *Listeria*. Pore-forming proteins also play a crucial role in immune defence, killing Gram-negative bacteria, cancer cells, and phagocytosed microbes^2^. Understanding how these proteins discriminate between self-cells and target membranes will provide insight into fundamental virulence strategies, as well as facilitate the application of engineered CDCs that lyse new cellular targets.

Many pore-forming proteins depend on lipid specificity to select their targets. For example, the immune protein perforin preferentially targets post-synaptic membranes through lipid disorder phases and neutral charge headgroups^3^. Another immune protein, gasdermin, specifically binds cardiolipin and phosphoinositide lipids to direct activity against mitochondria and the inner leaflets of eukaryotic plasma membranes, triggering cell death^4, 5^. Lipid specificity is a common theme for bacterial toxins in general^6^. For CDCs specifically, the requirement of cholesterol directs lytic activity towards plasma membranes of eukaryotic hosts^7^. Since cholesterol is not present in bacterial cells that secrete CDCs, this is one way the pathogen protects itself from damage during infection and toxin production; it also limits the repurposing of CDCs to attack bacterial pathogens. Many CDCs, such as pneumolysin (PLY), bind cholesterol directly through a conserved cholesterol-recognition motif located on a membrane-binding loop^8^. Although cholesterol is a well-known receptor for PLY, it may not be the only one. PLY also binds to a mannose receptor in dendritic cells, downregulating inflammation and promoting bacterial survival^9^. Other CDCs, such as intermedilysin (ILY), achieve species-specificity for their hosts by hijacking the cell surface receptor CD59 to initiate membrane-binding^10^. For this subgroup of CDCs, the interaction with CD59 is sufficient for attachment and the role of cholesterol is restricted to pore-formation^11^. While differences in membrane targeting may reflect diversity in CDC virulence strategies, lipid dependency for membrane penetration remains highly conserved.

CDCs bind target membranes through interactions located in their domain 4 (D4). Both the cholesterol recognition loops of PLY^8^ and CD59-binding site of ILY lie within this domain^10^. A long extended domain 2 (D2) flexibly links D4 to the membrane attack complex perforin (MACPF)/CDC (also referred to as D1 and D3) domain responsible for pore formation^12^. Membrane-binding of CDC monomers triggers a series of conformational re-arrangements that are required for oligomerization into an assembly referred to as a prepore. A dramatic vertical collapse of the oligomeric prepore brings two helical bundles (HB1 and HB2) within the MACPF/CDC domain close to the target membrane. HB1 and HB2 residues undergo rearrangement in secondary structure to form adjacent transmembrane β-strands in the pore^13-16^. These rearrangements are facilitated by stabilizing interactions between a number of amino acids at the MACPF/CDC domain interface^17^, but structural details of intermediate conformations remain unresolved.

Here we report the cryo-electron microscopy (cryoEM) structure of an early prepore-trapped conformation of the CDC intermedilysin (ILY). By binding to the human immune receptor CD59, ILY oligomerizes and triggers movement of a highly conserved amphipathic helix that encodes activity and lipid specificity of CDC pore formation. This helix can be modified to create cholesterol-independent CDCs or “super-CDCs” with significantly enhanced activity. Hence, our results provide a blueprint for engineering broad-purpose pore-forming proteins with tunable lytic activity.

## RESULTS

### CryoEM structure of an ILY early prepore

ILY from *Streptococcus intermedius* targets human cells by binding the GPI-anchored complement regulator CD59^10^ through an extended β-hairpin of D4^18^. To understand how this interaction initiates oligomerization on a target membrane, we trapped ILY in an early prepore state using a disulfide lock that restricts movement between D2 and the MACPF/CDC domain^11^. This disulfide-locked ILY variant binds CD59 and forms SDS-sensitive loosely-associated oligomers^11^, analogous to previously characterized CDC early prepore states^19^. We then used cryo-electron microscopy (cryoEM) to visualize this ILY variant in complex with CD59 anchored to cholesterol-containing lipid nanodiscs. We collected data on a Titan Krios microscope equipped with a direct electron detector and processed images using RELION^20^. 2D classification revealed heterogeneity in the number of subunits within a single nanodisc (Supplementary Fig. 1). Extensive 2D and 3D classification resulted in a final reconstruction comprised of 51,041 particles. The data were further refined with local symmetry, resulting in a map with an average resolution of 4.6 Å. (Supplementary Fig. 2). The final local resolution-sharpened density map illustrates an arc of five ILY-CD59 complexes, with the central subunit best resolved (Fig. 1a, Supplementary Fig. 1). We therefore built a model into the density of the central subunit and applied this fit as a rigid-body into the neighboring densities to model a three-subunit oligomer (Fig. 1a, Supplementary Fig. 3).

**Figure 1.**
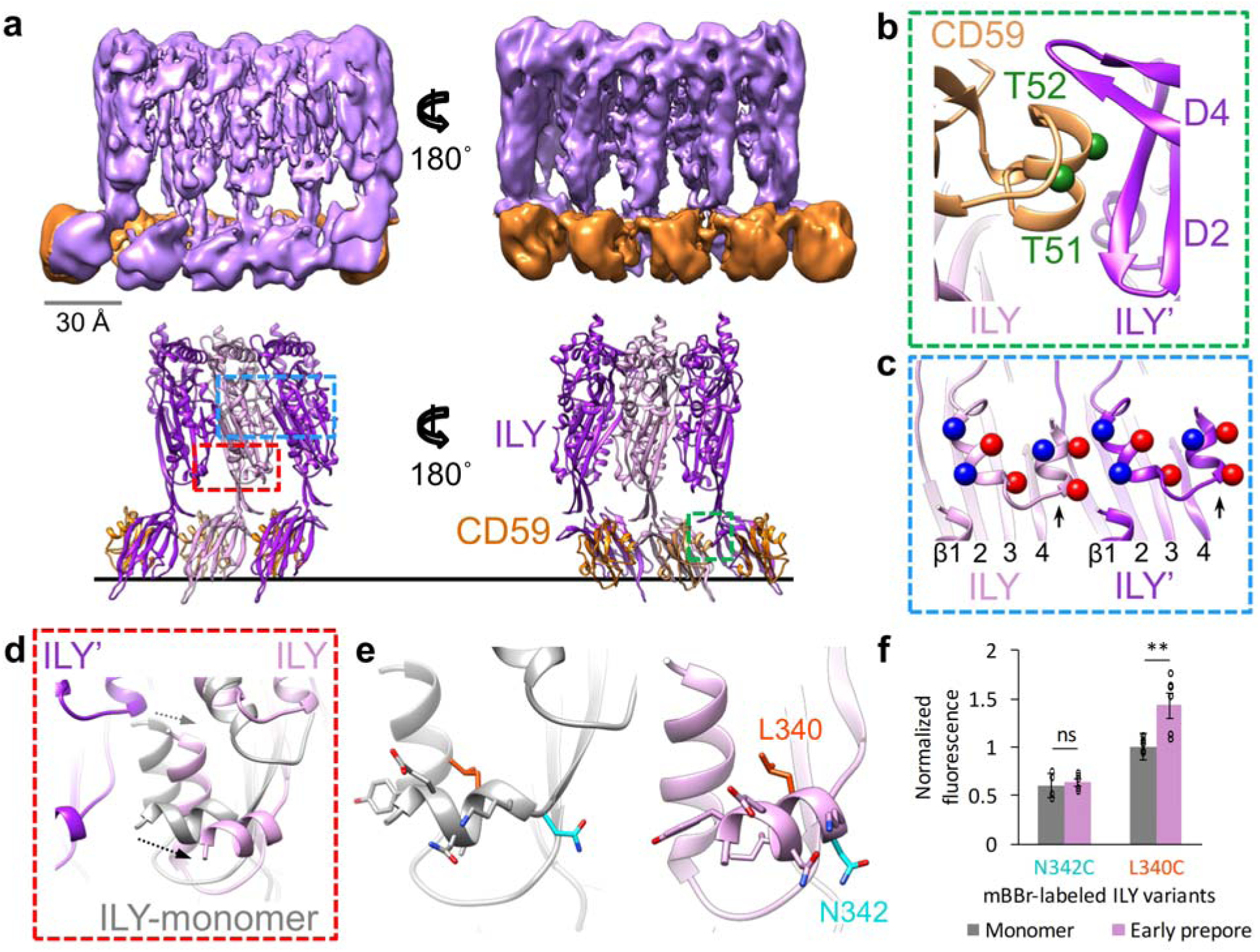
Structure of an ILY-CD59 early prepore oligomer. (a) CryoEM reconstruction filtered according to local resolution (top) and structural model derived from the cryoEM density map (bottom). In the model, adjacent ILY monomers (ILY and ILY’) are pink and purple, respectively. Neighboring CD59 molecules are orange. Black line represents the membrane surface. Dashed boxes indicate regions highlighted in panels b (green), c (blue), and d (red). (b) Interaction interface between CD59 and domains 2 and 4 (D2, D4) of neighboring ILY’. Green spheres indicate positions of residues modified by O-linked glycans in endogenous CD59 (T51, T52). (c) Helix-turn-helix motif of the oligomeric early prepore. Newly formed helix of ILY (formerly β-strand 5) is indicated by black arrow. Spheres indicate negative (red) and positive (blue) charged residues within the helices. β-strands of the MACPF/CDC domain are labeled. (d) Superposition of the early prepore ILY (pink) with monomeric ILY (grey) from crystal structure PDBID: 4BIK^18^. Grey and black arrows indicate direction of shifts for the vertical and horizontal helix (h-helix), respectively. Neighboring ILY’ is shown for reference. (e) Structural model showing the location of residues L340 (orange) and N342 (cyan) modified by the fluorescent tag, mBBr. (f) Normalized fluorescence intensity of mBBr-labeled h-helix residues in monomeric (grey) and oligomeric early prepore (pink) ILY. Individual data points shown as circles, n = 6, error bars represent standard deviation, p-value significance determined by student’s t-test: ns, not significant, p = 0.19; **, p = 0.0060. Sidechains have been added in COOT^42^ for visualization purposes.

**Figure 2.**
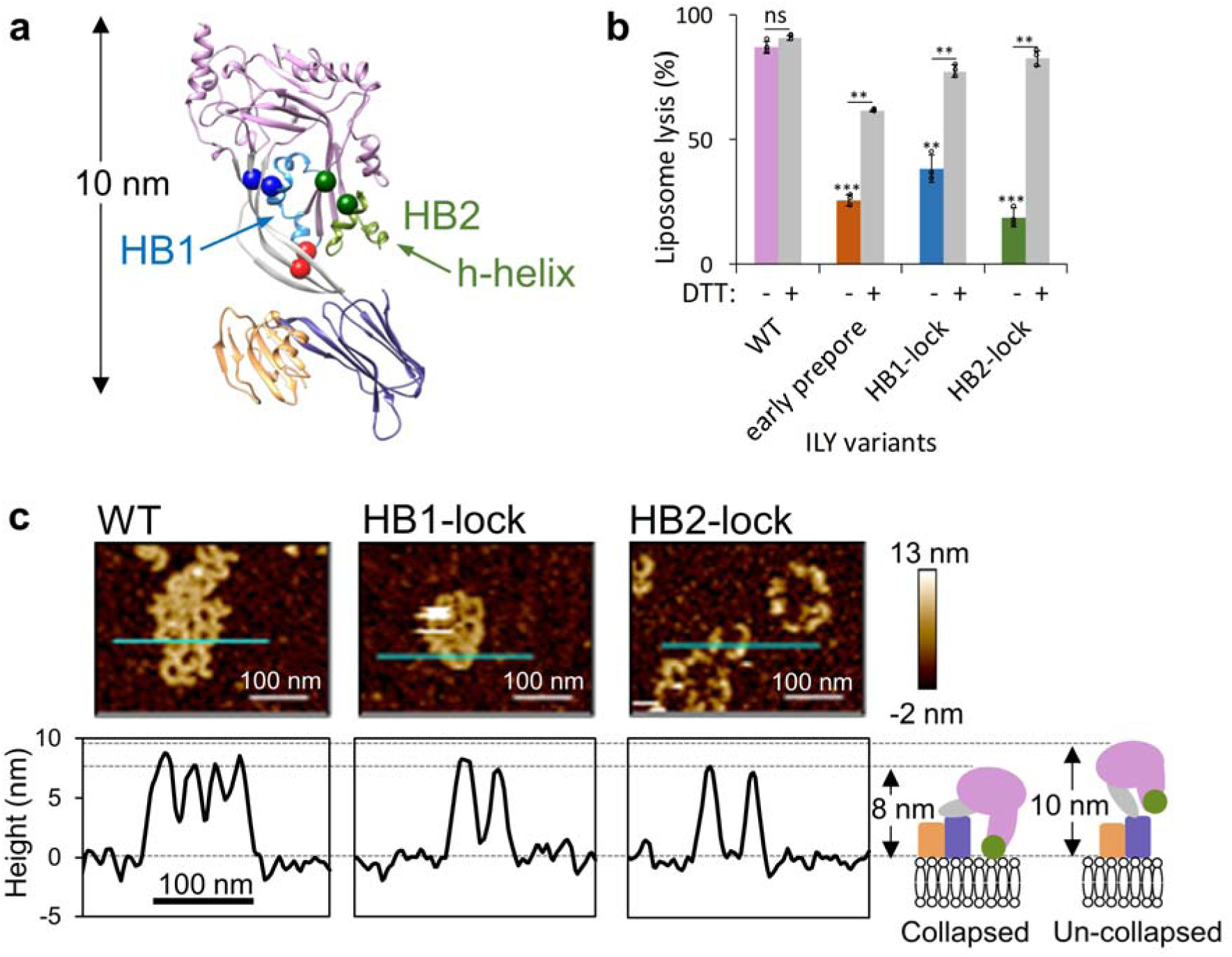
Trapping pore formation of collapsed ILY oligomers. (a) Early-prepore showing CD59 (orange) and ILY colored by domains: MACPF/CDC (pink), D2 (grey), D4 (purple). Domains 1 and 3 of ILY are grouped within the MACPF/CDC. HB1 (blue) and HB2 (green) are two helical bundles that form transmembrane β-hairpins in the final pore. Horizontal helix (h-helix) of HB2 is indicated (green arrow). Introduced cysteines (spheres) that form disulfide-bonds to trap conformations: early prepore (red), HB1-lock (blue) and HB2-lock (green). (b) A fluorescence-based calcein release assay was used to test lysis of liposomes containing cholesterol and CD59. ILY variants (wild-type (WT), early prepore, HB1-lock and HB2-lock) were analysed for activity with reducing agent (DTT) pretreatment. The statistical significance of this comparison is indicated above a horizontal line. Statistical significance displayed above each -DTT bar is for comparisons against wild-type ILY without reducing agent. Individual data points are circles, n = 3. Error bars represent standard deviation. P-value significance determined by 2-way ANOVA with a Bonferroni post-test: ns, not significant; **, p < 0.01; ***, p < 0.001. (c) AFM images (top) visualizing ILY variants on supported lipid bilayers containing cholesterol and CD59. Cyan line indicates image positions of height profiles (bottom) for ILY variants. Average assembly height and standard deviation was measured from 10 such profiles for each ILY variant: HB1-lock (7.9 nm +/- 0.6), HB2-lock (7.9 nm +/- 0.4), and WT (8.4 nm +/- 0.6). Schematic illustrates the domain organization (colored as in panel a) and vertical collapse measurements for ILY conformations.

**Figure 3.**
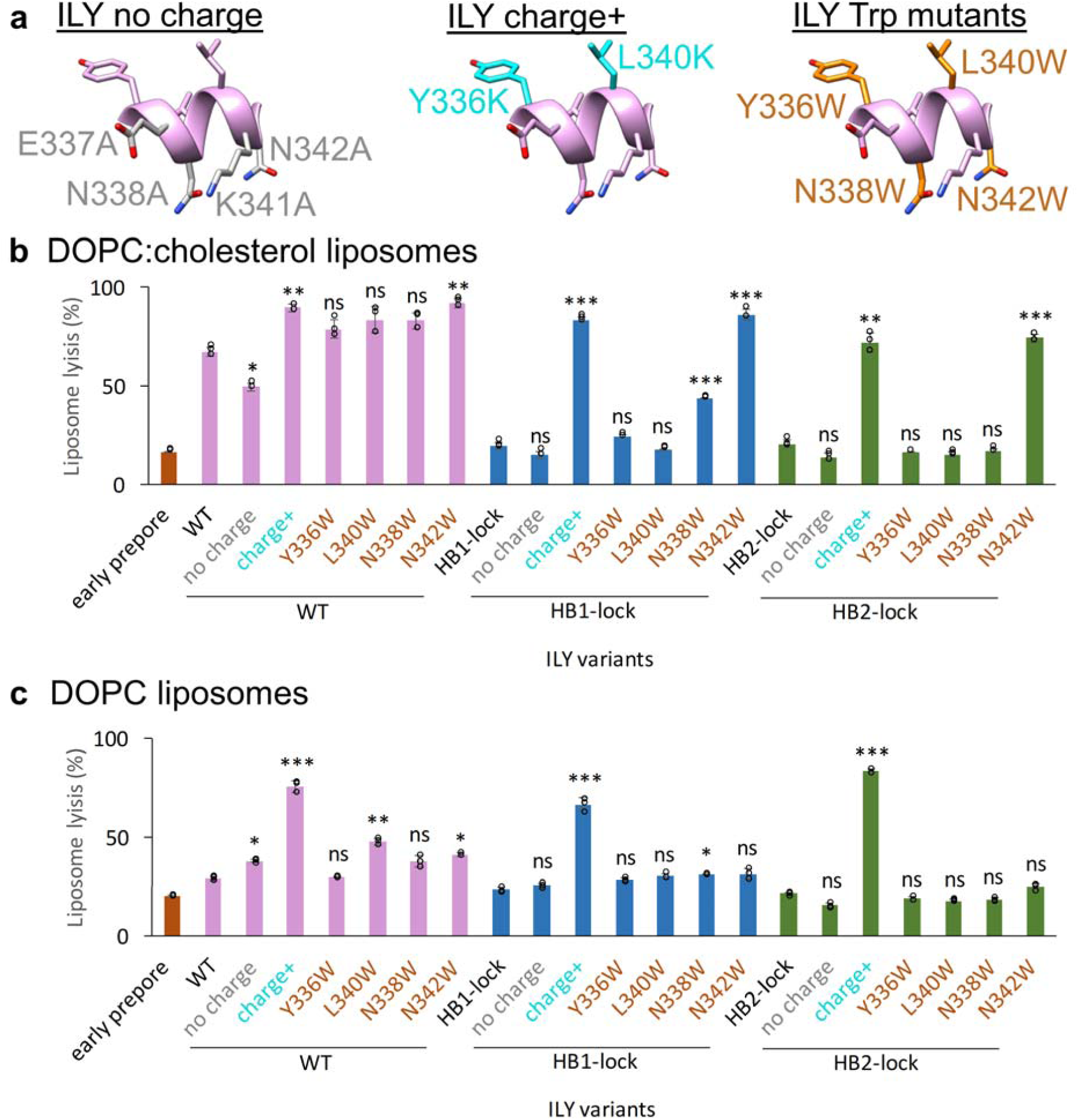
Targeted mutations in the horizontal helix (h-helix) tune lytic activity of ILY. (a) H-helix mutations are mapped onto the ILY structure. Left panel (ILY no charge) highlights residues (grey) exchanged for alanine (E337A, N338A, K341A, N342A) on the membrane proximal face of the h-helix. Middle panel (ILY charge+) indicates residues (cyan) mutated to lysine (Y336K, L340K) on the top face of the helix. Right panel (ILY Trp mutants) shows the location of residues (orange) mutated to tryptophan (Y336W, N338W, L340W, or N342W). Sidechains have been added in COOT^42^ for visualization purposes. (b, c) Activity of ILY h-helix mutants in wild type (WT), HB1-lock or HB2-lock backgrounds were tested using a calcein-release liposome lysis assay. Lysis of CD59-decorated liposomes comprised of DOPC:cholesterol lipids is shown in (b). Lysis of CD59-decorated DOPC liposomes is in (c). Statistical significance displayed above each bar is for comparison with the base ILY disulfide-locked variant (WT, HB1-lock, or HB2-lock, respectively). Individual data points shown as circles, n = 3. Error bars represent standard deviation. P-value significance determined by one-way ANOVA with a Bonferroni post-test: ns, not significant; *, p < 0.05; **, p < 0.01; ***, p < 0.01.

Our cryoEM structure reveals an oligomeric complex whereby a single CD59 can interact with two neighboring ILY monomers (ILY and ILY’). Similar to the soluble monomeric ILY-CD59 crystal structures^18, 21^, a β-hairpin of one ILY D4 extends the central β-sheet of CD59. The same β-hairpin of an adjacent monomer (ILY’), together with the tip of D2, sandwiches an α-helix of CD59 to form the second interface (Fig. 1b). This interface is dominated by two CD59 residues decorated with O-linked glycans in the native protein^22^. ILY binds to an O-glycan prevalent on human CD59^23^, suggesting that sugar recognition may play a role in oligomerization.

CD59-binding initiates the secondary structure re-arrangement of the MACPF/CDC outer β-strand (β-5). In our structure, residues within the β-5 strand form a new helix that comprises part of a helix-turn-helix motif (Fig. 1c). As a result, the β-4 strand is exposed to propagate oligomerization by binding the β-1 strand of a neighboring monomer. The ILY prepore assembly is most likely further stabilized by complementary charges on neighboring helical faces of the helix-turn-helix motif (Fig. 1c). Similarly, alternating charges of this motif in the PLY pore are thought to stabilize an inner barrel of helices above the membrane^16^.

In our complex, ILY extends to a vertical height of 110 Å, revealing an oligomeric prepore that has not yet collapsed towards the target membrane. Specifically, residues of the two MACPF/CDC helical bundles (HB1 and HB2) that eventually form transmembrane β-hairpins in the pore assembly do not contact the bilayer. Helical bundle 2 (HB2) is comprised of a vertical helix and a horizontal helix (h-helix) (Fig. 1d). Compared with the monomeric ILY-CD59, we observe a lateral shift in these helices upon oligomerization (Fig. 1d). This movement prevents an otherwise steric clash with a neighboring monomer and allows the oligomer to propagate. Given the moderate resolution of our map, we sought to verify the movement of these helices using a fluorescence-based assay. To investigate how oligomerization influences the local chemical environment of the h-helix, we covalently linked monobromobimane (mBBr) to residues on either the bottom (membrane adjacent) or top face of the h-helix in early prepore-locked ILY (Fig. 1e). Fluorescence was then measured in solution (monomer) or in the presence of CD59-containing DOPC:cholesterol liposomes (early prepore). Upon oligomerization, we observed an increase in mBBr fluorescence when the fluorophore was tethered to the top face of the h-helix, consistent with movement of L340 as it packs more closely against the surrounding protein. By contrast, the fluorescence intensity remained the same for labels attached to the bottom face of the h-helix, in agreement with our structural data showing that the packing of membrane-adjacent residues is not affected by this conformational change (Fig. 1e,f). Together, these biochemical data corroborate the structurally-observed lateral movement of the HB2 helices.

### The h-helix encodes the ability to tune lytic activity

Based on our cryoEM structure, we hypothesized that the h-helices of a collapsed oligomer could play a role in destabilizing the lipid bilayer. To test this, we first needed to stall pore formation of a collapsed oligomer by trapping ILY in a late prepore assembly intermediate. To this end, we generated disulfide-locked ILY mutants that restrict movement of either HB1 (HB1-lock) or HB2 (HB2-lock) and prevent the formation of transmembrane β-hairpins (Fig. 2a). Similarly to the ILY early prepore mutant, both HB1-lock and HB2-lock mutants show a significant reduction in lytic activity, which is rescued under reducing conditions (Fig. 2b). These data, together with a low level of free cysteines (Supplementary Fig. 3), confirm that the engineered disulfide bonds obstruct pore formation of HB1-lock and HB2-lock mutants. To investigate if these ILY variants formed vertically collapsed oligomers, we imaged complexes using atomic force microscopy on supported lipid bilayers containing both cholesterol and CD59. In contrast to monomeric ILY^11^ and the ILY early prepore (Fig. 2a), which extend 10 nm from the membrane surface, both HB1-lock and HB2-lock mutants form collapsed oligomers analogous to the 8 nm high wild-type pores (Fig. 2c), yet show significantly reduced lytic activity (Fig. 2b). We conclude that HB1-lock and HB2-lock mutants trap collapsed prepore assemblies whereby the helical bundles are brought closer to the target membrane but do not breach the bilayer.

The h-helix is amphipathic, with charged or polar residues along the membrane-adjacent face and hydrophobic residues on the opposite face (Fig. 3a). To test how lytic activity and hence ILY function depend on the distribution of charges along the ILY h-helix, we generated ILY variants where charged or polar residues on the membrane proximal face were substituted with alanine (ILY^no charge^) or where hydrophobic residues on the top face were swapped with lysine (ILY^charge+^) (Fig. 3a). Removal of charged residues impaired lytic activity of wild-type ILY. By contrast, incorporating positively charged amino acids on the top face of the h-helix increases lysis. Lytic activity of wild type ILY could also be improved by substituting h-helix residue N342 for tryptophan (Fig. 3b). Taken together, these results demonstrate that the lytic function of ILY can be modulated by changing the physiochemical properties of the h-helix.

By introducing mutations in the HB1-lock and HB2-lock backgrounds, we next analyzed the effect of h-helix variations on otherwise dysfunctional (non-lytic) collapsed prepores. Remarkably, we found that the lytic function was fully recovered by the addition of charged residues (charge+) or the N342W substitution in the h-helix, as shown on cholesterol-containing liposomes (Fig. 3b). By contrast, tryptophan substitutions elsewhere along the helix only partially recovered lytic activity for the HB1-lock mutant and did not enhance lysis at all for the HB2-lock mutant (Fig. 3b). We deduce that tryptophan substitutions along the bottom face of the h-helix augment lysis by intercalating into the outer leaflet of the bilayer and anchoring the h-helix to the membrane. The position of the tryptophan likely affects the efficiency of this mechanism, since substitution at N342 shows a stronger phenotypic change than at N338.

Since ILY only requires cholesterol to rupture the membrane and not to bind it^11^, the ILY-CD59 system offers an opportunity to explicitly investigate and potentially modulate the cholesterol-dependency of CDC membrane lysis. We therefore measured the lytic activity of our ILY variants on liposomes without cholesterol (Fig. 3c). Notably, the requirement of cholesterol was abrogated by the addition of positively charged residues on the top face of the h-helix (charge+); this modification resulted in high levels of lysis in all ILY backgrounds (wild-type, HB1-lock, and HB2-lock). When the charges are removed from the h-helix of wildtype ILY, we found modest, but opposite changes in lysis of the two types of liposomes tested here (Fig. 3b,c), and no significant effect was observed in the HB1-lock or HB2-lock backgrounds. Furthermore, compared with the results on cholesterol-containing liposomes, N342W substitution in HB1-lock and HB2-lock backgrounds did not yield a similarly dramatic increase in lytic activity. Together, these data demonstrate that modifications of the h-helix can tune lytic activity in different lipid environments.

The amphipathic nature of the ILY h-helix is highly conserved across CDCs. Structures of both CD59-dependent (vaginolysin: VLY) and CD59-independent CDCs (pneumolysin: PLY, suilysin: SLY, perfringolysin: PFO, and listeriolysin O: LLO) contain a similarly oriented amphipathic helix within HB2 (Fig. 4a). To test if the principles of tuning lytic activity can be extended to other CDCs, we introduced similar charge-altering mutations in PLY. As PLY requires cholesterol to initiate membrane attachment, we only used cholesterol-containing liposomes to compare lytic activity of PLY mutants with respect to wild-type PLY. In agreement with our results for ILY, PLY lytic activity was decreased when charged and polar amino acids of the h-helix were substituted with alanine (PLY^no charge^). Moreover, the lytic activity was enhanced by replacing uncharged residues on the top helical face with lysines (PLY^charge+^), again analogous to what we observed for ILY variants (Fig. 4b,c). These data suggest that modifications within the h-helix provide a generic mechanism for controlling CDC pore formation and lytic activity.

**Figure 4.**
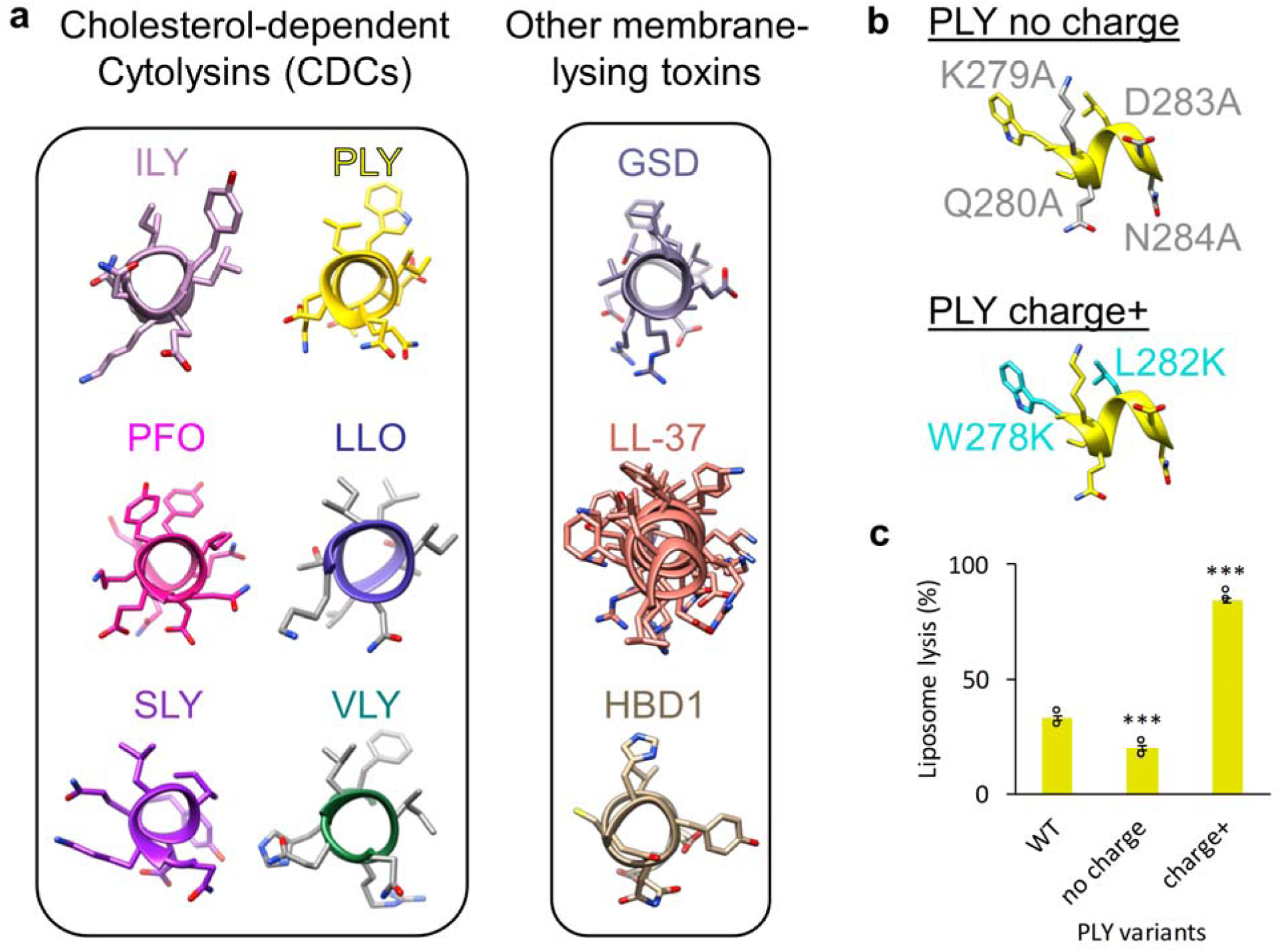
H-helix modifications tune activity of a non-CD59 binding CDC. (a) The equivalent h-helix in cholesterol-dependent cytolysins (CDCs) are shown for intermedilysin (ILY, PDBID: 4BIK)^18^, pneumolysin (PLY, PDBID: 5AOE)^16^, perfringolysin O (PFO, PDBID: 1M3I)^47^, listeriolysin O (LLO, PDBID: 4CDB)^48^, suilysin (SLY, PDBID: 3HVN)^49^, and vaginolysin (VLY, PDBID: 5IMY)^21^. Amphipathic membrane-interacting helices are shown for other membrane-lysing proteins: gasdermin (GSD, PDBID: 6CB8)^28^, the CAMP cathelicidin (LL-37, PDBID: 5NMN)^50^, human beta-defensin 1 (HBD1, PDBID: 1IJV)^51^. (b) H-helix mutations in PLY analogous to the ILY variants are mapped onto the PLY structure (PDBID: 5AOE). The positions of alanine substitutions (K279A, Q280A, D283A, N284A) on the PLY h-helix are shown in grey (no charge). Residues on the PLY h-helix mutated to lysine (W278K, L282K) are highlighted in cyan (charge+) (c) The ability of PLY h-helix variants to lyse cholesterol-containing liposomes was assessed using a calcein-based lysis assay. Statistical significance displayed above each bar is for comparisons with wild-type PLY (WT). Individual data points shown as circles, n = 3. Error bars represent standard deviation. P-value significance determined by one-way ANOVA with a Bonferroni post-test: ***, p < 0.001.

## DISCUSSION

Our combined structural and biochemical data support a model whereby CDC oligomerization triggers movement of an amphipathic helix that modulates lytic activity (Fig. 5). Soluble CDC monomers associate with membranes through interactions within D4. Binding is mediated either by loops at the tip of D4 with cholesterol^8^, or by engaging cell surface receptors, such as CD59^10^. Membrane-binding initiates oligomerization and conformational changes within the MACPF/CDC domain that create new interaction interfaces between monomers. Specifically, the formation of a helix-turn-helix motif exposes the MACPF/CDC β-4 strand to propagate the oligomer; it may also contribute to stability of the assembly. In our structure of an early prepore, the h-helices of ILY are arranged parallel to the membrane with their charged surfaces facing the outer leaflet of the lipid bilayer. Following a dramatic vertical collapse of the oligomeric prepore, the h-helix is brought in close proximity to the target membrane. Lying along the surface, the amphipathic h-helices of the oligomer would then cause local membrane disruption, analogous to the ‘carpet’ model of membrane disruption by cationic antimicrobial peptide (CAMP)^24^ amphipathic helices (Fig. 4a). The helix-to-hairpin transition of CDC transmembrane residues is likely to occur via several intermediate structural states^25^ with different local energy minima, including collapsed coil-like intermediates^26^. The transient states of the h-helix may contribute to disruptive forces that prime the membrane for hairpin insertion and to the displacement of lipids by water molecules at the inner hydrophilic surface of the pore, as observed by molecular dynamics simulations^27^. We demonstrate that modifications of the h-helix similarly modulate the lytic activity of a non-CD59 binding CDC, PLY. Furthermore, amphipathic helices are present in and influence the lytic activity of other oligomeric pore-forming proteins such as gasdermin^28^ and MPEG-1^29, 30^. Taken together, our model may reveal a fundamental mechanism underpinning membrane rupture by oligomeric pore-forming proteins.

**Figure 5.**
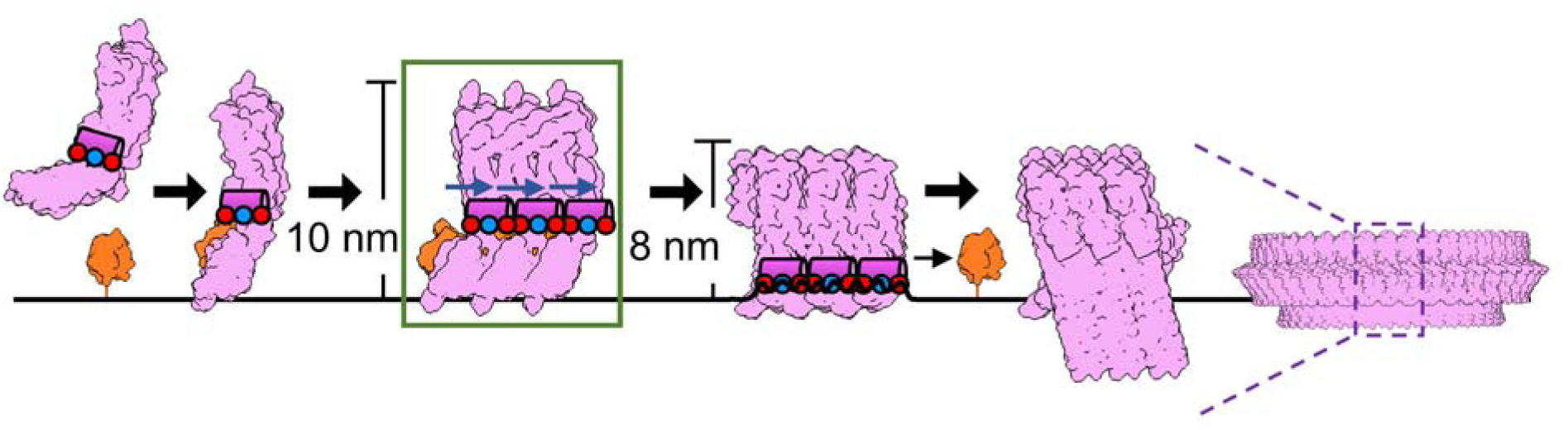
Model for CDC pore formation and membrane lysis. Soluble monomers (pink) bind target membranes through interactions with either cholesterol or cell surface receptors, such as CD59 (orange). Oligomerization of the membrane-bound subunits (green box) causes structural re-arrangements that include a shift (blue arrow) of the amphipathic h-helix (pink cylinder), as captured in our cryoEM structure. The early prepore, which extends 10 nm from the lipid bilayer, collapses to 8 nm and brings the charged face of the h-helix (blue and red circles) into contact with the membrane. The amphipathic h-helix disrupts the membrane, leading to the transition of helical bundles (HB1 and HB2) into membrane-piercing β-hairpins. For ILY, CD59 dissociates, as it is not part of the final pore. Though only three monomers are displayed, this process occurs for higher oligomer arc and ring-like pores^15^. Figure is based on ILY and PLY structures: PDBIDs 1S3R^52^, 4BIK^18^, 5CR6^16^, and 5LY6^16^; and EMD-4118^16^.

Our results open an exciting perspective for protein engineering of h-helices for the creation of “super-CDCs” with significantly enhanced lytic activity or of cholesterol-independent CDCs that can be used to target bacterial pathogens. Specifically, our results show that CDC-mediated lysis is enhanced by the addition of positive charges to the top face of the h-helix, and that this removes the requirement of cholesterol for membrane rupture. We note that the tunable lysis of the h-helix bears similarities to CAMP-mediated cell lysis, where the membrane specificity is encoded by the charge properties of the helix^31^; and where the introduction of tryptophan residues can modulate antimicrobial activity of synthetic CAMPs^32^. By combining h-helix modifications with mutations in membrane binding regions of D4^9, 33, 34^, CDCs could be designed to lyse new cellular targets such as plant-pathogenic bacteria. The devastating effects of these pathogens on crop yield and productivity^35^ could be eliminated by engineered pore-forming proteins that lyse bacterial membranes but are unable to penetrate a plant cell wall.

## ONLINE METHODS

### Bacterial strains and plasmids

*Escherichia coli* strain DH5α was used to maintain plasmids and for cloning purposes. Strains BL21 DE3 and BL21 DE3 pLysS were used for expression of ILY and PLY variants, with the latter strain grown in the presence of 25 µg/ml chloramphenicol to maintain the pLysS plasmid. His-tagged ILY and PLY constructs were cloned and expressed using pTricHisA and maintained with 100 µg/ml ampicillin. His-tagged MSP2N2^36^ was expressed from pET28-MSP2N2 (Addgene) in the presence of 50 µg/ml kanamycin.

### Generation of ILY and PLY variant constructs

Wild-type ILY and PLY, for this study, have been codon optimized for *E. coli* and all native Cys have been mutated to Ser^11^. All ILY and PLY variants were created by site-directed mutagenesis, in which a pTricHisA plasmid with the template gene was amplified with pairs of primers that contain the desired mutations (Supplementary Table 1) and Q5 DNA polymerase (New England Biolabs). This was followed by DpnI (New England Biolabs) digestion, to remove the template plasmid, heat-shock transformation into *E. coli* DH5α, and verification by sequencing. CDC variants used in this study are summarised in Supplementary Table 2.

### Purification of ILY and PLY variants

*E. coli* BL21 DE3 containing pTricHisA with an *ily* or *ply* variant gene were grown to an OD_600_ of ∼0.6-0.9 in Luria-Bertani (LB) broth at 37°C, and induced with a final concentration of 50 µg/ml of isopropyl β-d-1-thiogalactopyranoside (IPTG) at 18°C for 18 hours. In the case of ILY-WT^charge+^ and ILY-HB2lock^charge+^, this method resulted in very low levels of protein expression. Therefore, these two ILY variants were expressed in *E. coli* BL21 pLysS and induced with 50 µg/ml IPTG at 18°C for 1 hour. The cells were collected by centrifugation and lysed by sonication in Buffer A (200 mM NaCl, 20 mM Tris-HCl, pH 7.5) containing cOmplete Protease Inhibitors (Roche). The soluble fraction was incubated with HisPur Cobalt resin (Thermo Fisher Scientific), washed with 10 mM imidazole in Buffer A. Bound His-tagged proteins were eluted with 100 mM and 500 mM imidazole in Buffer A. His-tagged protein was further purified by size exclusion chromatography over a Superdex 200 10/300 column (GE Healthcare) in Buffer A.

### Forming the ILY early-prepore on CD59-decorated lipid nanodiscs

MSP2N2 protein was first produced by growing BL21 DE3 cells containing pET28-MSP2N2 (Addgene) to an OD_600_ of ∼0.8 in LB and inducing with 0.5 mM IPTG at 37°C for 2-3 hours. Cells were pelleted and reconstituted in 40 mM Tris-HCl pH 7.4 with cOmplete Protease Inhibitors (Roche), to which TritonX-100 was added to a final concentration of 1%. Cells were stored on ice for 10 min, then lysed by sonication and supplemented with 0.3 M NaCl. His-tagged protein was bound to HisPur Cobalt resin (Thermo Fisher Scientific), washed with 40 mM Tris-HCl pH 8.0, 0.3 M NaCl, 0.5% Triton X-100, then washed with 40 mM Tris-HCl pH 8.0, 0.3 M NaCl, 50 mM sodium cholate. Protein was then eluted into 40 mM Tris-HCl pH 8.0, 0.1 M NaCl, 300 mM imidazole and subjected to size exclusion chromatography over a Superdex 200 10/300 column (GE Healthcare) with 40 mM Tris-HCl pH 8.0, 0.1 M NaCl, 0.5 mM EDTA. Nanodiscs were created by mixing 3.44 mM MSP2N2 with 34.4 mM lipids (DOPC:cholesterol, 1:1 molar ratio) solubilized in 40 mM Tris-HCl pH 8.0, 100 mM NaCl, 0.5 mM EDTA, 64 mM sodium cholate, such that the final sodium cholate concentration was 32.5 mM. This mixture was then incubated on ice for 30 minutes followed by an overnight incubation with activated Bio-beads SM2 (Bio-Rad) at 4°C with agitation, to remove the detergent and promote nanodisc formation. The nanodiscs were then purified by size exclusion chromatography with 40 mM Tris-HCl pH 8.0, 0.1 M NaCl, 0.5 mM EDTA over a Superose 6 10/300 column (GE Healthcare) and stored at 4°C.

A recombinant extracellular domain of CD59 modified with a lipid-anchoring peptide was gifted by R.A.G. Smith (King’s College London). Briefly, the cytoplasmic domain of human CD59 modified with an additional C-terminal cysteine was expressed in *E*.*coli* and purified from inclusion bodies^37^. The cytotopic modification reagent bis-myristoyl lysyl SSKKSPSKKDDKKPGD (S-2-thiopyridyl)-cysteine acid (APT3146, Cambridge Research Biochemicals) was covalently linked to the C-terminal cysteine and the modified protein was purified by hydrophobic interaction chromatography and ammonium sulfate precipitation^18, 38^.

MSP2N2 nanodiscs were incubated with 3 µg/ml CD59 at room temperature for 20 minutes in Buffer A, followed by another 20-minute incubation after the addition of 20 µg/ml ILY-prepore. The mixture was immediately washed twice with Buffer A on an Amicon Ultra 0.5 ml 100 kDa concentrator column (Merck) to remove unbound protein and concentrated by a factor of 5. To avoid particle aggregation, the final sample was immediately used to prepare EM grids.

### Negative stain EM

All EM samples were first screened by negative-stain EM, to assess particle integrity and distribution. 2.5 µl of ILY-prepore-CD59 on nanodiscs was applied to glow-discharged, carbon-coated copper grids (Agar Scientific) and stained with 2% uranyl acetate. Samples were imaged on a 120 keV Tecnai T12 microscope (Thermo Fisher Scientific) with a 2K eagle camera (FEI) at a nominal magnification of 50,000x for evaluation.

### CryoEM grid preparation and data collection

ILY-CD59 nanodisc complexes were imaged using holey carbon grids coated with graphene oxide. To coat R1.2/1.3 Quantifoil grids with graphene oxide, grids were first glow-discharged for 1 min, then, 0.2 mg/ml of a graphene oxide solution (Sigma) in water was applied to the glow-discharged, top face of the grid, followed by blotting by filter paper on the bottom face of the grid. This process was repeated twice, followed by two washes of the top face of the grid with 20 µl of water. Grids were left to dry, and used within one hour of graphene oxide coating. Immediately following concentration, 2.5 µl of the early prepore-locked ILY-CD59 oligomers on nanodiscs was adsorbed on graphene oxide-coated grids and blotted for 2.5 seconds at ‘blot force’ 3 and plunge frozen in liquid ethane cooled to liquid nitrogen temperatures with a Vitrobot mark III (Thermo Fisher Scientific). Electron micrograph movies were collected on a 300 keV Titan Krios (Thermo Fisher Scientific) fitted with a Falcon III direct electron detector (Thermo Fisher Scientific) in linear mode with image acquisition software EPU (Thermo Fisher scientific). Specific collection details for all 3 data sets are summarized in Supplementary Table 3.

### Initial model generation

An initial model was generated in RELION^20^ from a dataset collected on a 300 keV Titan Krios (Thermo Fisher Scientific) with a Quantum K2 Summit direct electron detector (Gatan) in counting mode with a magnified pixel size of 1.048 Å. Manually picked particles from motion corrected and ctf-estimated micrographs were subjected to rounds of 2D classification and class curation, resulting in 4241 particles which were used to generate an initial model in RELION^20^. 3D refinement with this first initial model produced a reconstruction with an average resolution of 8.6 Å. This density was then used as the initial model for the first individual 3D reconstructions of data sets 1, 2, and 3. Particles used to generate the initial model were not included in the refinement of subsequently collected datasets 1, 2, and 3.

### CryoEM data analysis and density reconstruction

The overall data analysis and reconstruction strategy is summarized in Supplementary Figure 3. All data analysis and reconstruction were completed via RELION^20^ unless otherwise stated. Data sets 1, 2, and 3 were treated separately for the initial stages of data processing. For each data set, micrograph movie frames were aligned using MotionCor2^39^ and CTF parameters were estimated with CTFFIND4^40^. Any flattened movies containing low figure of merit scores, crystalline ice, low contrast, or substantial drift were removed from further analysis. Particles were picked with a combination of manual picking, autopicking in RELION, and crYOLO^41^, and duplicate particles were removed. Initial 2D class averages included signal from the nanodisc (Supplementary Fig. 1). To improve the alignment of the particles based on features of the ILY oligomer, particles were re-centred on the early prepore signal with a smaller mask to exclude the nanodisc. After several rounds of 2D classification and class selection, the remaining particles were reconstructed into 3D density using the initial model low pass filtered to 40 Å. The particles from data set 3 were collected at a 30° tilt to improve angular distribution of the particles and were therefore subjected to per-particle CTF refinement, followed by another round of auto-refinement. At this stage all particles from data sets 1, 2, and 3 were pooled, and after a final 2D classification, 105,448 particles were selected for 3D density reconstruction, resulting in an average resolution of 4.8 Å. To improve homogeneity in the particles, they were classified into 4 groups with local angular searches performed during each iteration. The combination of 2 classes (51,041 particles) produced the best quality density reconstruction, with an average resolution of 4.7 Å. Next, local symmetry was imposed during the 3D reconstruction, with local angular searches. Two strategies were used in an attempt to improve the density of different regions. First, the top region (ILY MACPF/CDC domain and D2) of two monomers were designated as locally symmetric during 3D reconstruction (Supplementary Fig. 3, bottom right branch). Secondly, in an attempt to further optimize the density of the bottom region (ILY D4 and CD59), both the top and bottom regions of two monomers (Supplementary Fig. 3, bottom left branch) were specified in local symmetry operators during refinement. After this, each reconstruction was refined with the same local symmetry designations, but in this case, references were low pass filtered 10 Å and only local searches were performed. The estimated resolution for the density map in which the top regions were symmetrized is 4.6 Å (Supplementary Fig. 1). The final density map for this reconstruction was generated by local resolution filtering in RELION, with a global B-factor of -220. This reconstruction had the best local resolution estimates and density features over the entirety of the middle monomer, including the bottom region (Supplementary Fig. 2, comparing bottom left and right branches), and was therefore used for building and refining the structural model. Though the average resolution estimated for each of these steps only improved from 4.8 to 4.6 Å, there was a noticeable increase in the quality of the sharpened local resolution-filtered map after each stage of processing.

### Structural model building and refinement

The crystal structure for soluble, monomer-locked ILY (PDBID 4BIK^18^) was placed in the middle monomer of the local resolution-filtered density with COOT^42^. The new helix-turn-helix motif in the MACPF/CDC domain was built using the pneumolysin pore structure as a reference (PDBID 5LY6^16^). The vertical and horizontal helices of HB2 were shifted into the density with the real space refine tool in COOT^42^. Side chains were then removed in COOT before refinement with Phenix real_space_refine. The model was initially refined as 3 rigid bodies. The MACPF/CDC domain was segmented into domains: D1 and D3 as is convention. The first body comprised CD59 and ILY D4; the second was made up of D1 and D2; the third body was D3. The helix-turn-helix motif and HB2 helices were further refined with global minimization and secondary structure restraint to generate the final model in Phenix^43^. Coordinates were also independently refined with Namdinator, which combines Phenix real_space_refine with molecular dynamics simulations^44^. Both refinements produced consistent models of the h-helix and HB2 helices (RMSD of alpha carbons: helix-turn-helix, 1.55 Å; domain 3 HB2, 1.50 Å). To generate the oligomeric structure of three ILY-CD59 subunits, the coordinates of the central monomer (Supplementary Fig. 3) were rigid body fit into the neighboring densities using COOT^42^ and Phenix real_space_refine^43^. Models were validated with the Phenix cryoEM validation tool^45^ (Supplementary Table 4). Structures are deposited to EMDB and PDB under accession numbers: EMD 11172 and PDB 6ZD0.

### Fluorescence quenching assay

Fluorescently labeled ILY variants were generated by mutating L340 or N342 to a cysteine residue and covalent modification with monobromobimane (mBBr). ILY variants were incubated with mBBr at 10°C overnight under agitation with a 1:5 molar ratio of ILY:mBBr. The free dye was removed by buffer exchange in Zeba spin 0.5 ml columns (Thermo Fisher Scientific) and mBBr-labeled protein was stored in the dark at -80°C to preserve the fluorescent dye.

To test fluorescence of soluble, monomeric ILY, 9 µg/ml ILY-prepore-mBBr was mixed with 1.5 µg/ml CD59 in Buffer A; and for the oligomeric, membrane-bound condition, the ILY-prepore-mBBr and CD59 mixture also included 0.375 µg/ml of DOPC:cholesterol (1:1 molar ratio) liposomes^6^. Fluorescence was measured by excitation at 398 nm, and an emission spectrum was collected from 430-600 nm on a CLARIOstar plate reader (BMG labtech), with the spectrum peak taken as the emission fluorescence reading. Fluorescence of each sample was normalized to the fluorescence of denatured protein (addition of 10% SDS) by dividing the monomer or prepore fluorescence by the denatured fluorescence value of each respective sample.

### Liposome lysis assay

Lytic activity of CDC variants was assessed using a calcein-release liposome lysis assay. Liposomes containing calcein were first prepared by rehydrating lipids (DOPC or DOPC:cholesterol, 1:1 molar ratio) in Buffer A with 50 mM calcein, followed by size exclusion chromatography with Sephadex G-50 resin (Sigma) in Buffer A with 500 mM sucrose, as previously described^11^. The resultant liposomes were filled with self-quenching calcein dye while external, unquenched dye was replaced by Buffer A and sucrose. To determine the activity of ILY variants, liposomes were first incubated with a final concentration of 1.0 µg/ml CD59 for 20 minutes at room temperature, followed by the addition of ILY at a final concentration of 9.0 µg/ml. The activity of PLY variants was determined by adding a final concentration of 48.4 µg/ml of PLY to DOPC:cholesterol liposomes. Fluorescence intensity was read with CLARIOstar plate reader (BMG labtech) at an excitation wavelength of 490 nm and emission wavelength of 520 nm. Total liposomes lysis was achieved by the addition of 0.87% n-dodecyl-beta-maltoside (DDM) and a freeze/thaw cycle at -80°C. To calculate percent lysis, the fluorescence value from a buffer well was first subtracted from all raw fluorescence readings at a single time point between 30 and 60 minutes. Each blanked-corrected experimental reading was divided by the corresponding total liposome lysis fluorescence reading (also blank-corrected). Activity of disulfide-locked CDC variants was assessed by preincubating the protein with or without 20 mM DTT for 30 min at room temperature.

### Cysteine-accessibility assay

The Protein Thiol Fluorescent Detection kit (Invitrogen) was used to determine the percent of free Cys in disulfide-locked ILY variants. After incubation with the detection reagent, fluorescence intensity was read with a CLARIOstar plate reader (BMG labtech) with an excitation wavelength of 390 nm and emission wavelength of 510 nm to measure the level of free Cys. The fluorescence measurement was then converted to a concentration of free thiols from a standard curve. The percentage of free Cys was calculated from the total moles of Cys present in each sample.

### Atomic force microscopy sample preparation

For the preparation of supported lipid bilayers, pure lipids were dissolved in chloroform at 10 mg/mL and mixed in solution to give a lipid mixture of DOPC:cholesterol 2:1 molar ratio. The lipid-in-chloroform solution was then dried in a glass vial under a stream of nitrogen gas to give 1 mg of lipid as a thin film. The lipid film was hydrated in buffer (20 mM Tris, 200 mM NaCl, pH 7.5), vortexed and bath sonicated to give a cloudy lipid suspension. The suspension was then passed through a 50 nm polycarbonate membrane (GE Healthcare Lifesciences) 15 times to yield a clear suspension of small unilamellar vesicles.

Supported lipid bilayers were formed by injecting 4.5 µL of the lipid vesicle suspension to a freshly cleaved mica disk (6 mm diameter) under 18 µL of incubation buffer (hydration buffer plus 10 mM CaCl_2_ solution at 37°C; this induces the rupture of the vesicles onto the mica support over an incubation period of approximately 30 minutes. Excess vesicles were then removed from the supernatant by rinsing with 500 µL of the hydration buffer, to yield a uniform bilayer free of adsorbed vesicles (as assessed by AFM imaging). The supported lipid bilayers were next incubated with a final concentration of 50 ng/ml CD59 for 5 minutes, and thereafter with a final concentration of 100 µg/ml ILY for 15 minutes, all at 37°C, and next washed with 500 µL of the hydration buffer.

### Atomic force microscopy imaging and analysis

AFM imaging was performed in buffer solution using a Multimode 8 AFM (Bruker, Santa Barbara, USA) and MSNL-E and PEAKFORCE-HIRS-F-B cantilevers (Bruker), in off-resonance tapping / fast force-feedback imaging (Bruker’s PeakForce Tapping) mode where force-distance curves were recorded at 2 kHz, with amplitudes of 10-20 nm. The experiments were performed at room temperature, largely following procedures previously described elsewhere^3^.

We note that (membrane-inserted) pore assemblies are readily imaged by AFM, as they make contact, through the membrane, with the underlying solid support; but that prepore assemblies are rather hard to image by AFM, as the supported lipid bilayers generally represent too fluid a support to retain the assemblies in place^3^, except when the assemblies form larger clusters. In this case, the mobility of HB1/HB2-lock prepore assemblies may also be constraint by the dense coverage of CD59 on the membrane surface^11^.

Images were processed using open-source SPM analysis software, Gwyddion (v2.55)^46^ for first-order plane-fit background subtraction and second-order line-by-line flattening. The membrane surface was referenced as zero height. The images in Fig. 2c were cropped from 500 x 500 nm^2^. Assembly heights (mean ± standard deviation) were estimated based on 10 height profiles for 10 different assemblies from each 500 x 500 nm^2^ image, referenced with respect to the membrane surface, as exemplified in Fig. 2c.

### Statistical analysis

The student’s t-test (assuming equal variance) was used to assess statistical significance of differences between monomer and early prepore fluorescence quenching of h-helix-labeled ILY (ILY-prepore^N342C^ or ILY-prepore^L340C^). To compare the lytic activity of disulfide-locked mutants in the presence or absence of DTT, first a two-factor ANOVA test was performed to confirm significant variance amongst the samples, followed by a Bonferroni post-test for paired comparisons. A single-factor ANOVA test was used to determine the significance of variance within datasets describing lytic activity of h-helix mutants (ILY and PLY). A Bonferroni post-test was used for paired comparisons within each set of h-helix variants (ILY-WT, ILY-HB1lock, ILY-HB2lock, PLY).

## ACKNOWLEDGEMENTS

We thank R. Smith for gifting the CD59; Y. Chaban for data acquisition assistance; S. Islam for computational support; J. Demmer for assistance with model building; P. Haynes for guidance on the AFM image analysis; and A. Menny and the Bubeck lab for discussions. We thank Diamond for access and support of the Cryo-EM facilities at the UK national electron bio-imaging centre (eBIC), proposal EM18659, funded by the Wellcome Trust, MRC, and BBSRC. D.B. and N.R.S. are supported by a CRUK Career Establishment Award (C26409/A16099) to D.B.; C.M.B. and T.B.V. are funded by BBSRC Doctoral Training Program grants, Ref: BB/J014575/1 and BB/M011178/1, respectively. E.S.P. and B.W.H. have been supported by EPSRC and MRC (EP/M507970/1 to E.S.P.; MR/R000328/1 to B.W.H.), and acknowledge EPSRC investment in AFM equipment (EP/M028100/1).

## AUTHOR CONTRIBUTIONS

N.R.S. conducted cryoEM work, generated ILY mutants and performed ILY fluorescence and lysis experiments. N.R.S. and D.B. conceived the ideas, analyzed the results, and wrote the manuscript. T.B.V. generated PLY mutants and ILY-prepore^charge+^, conducted PLY and ILY-prepore^charge+^ lysis experiments, and analyzed these data. T.B.V., E.S.P., and B.W.H. analyzed AFM data. E.S.P. performed AFM experiments. C.M.B. generated the ILY-HB1lock mutant. All authors assisted with manuscript editing.

