## Supplementary_material for "Structural basis for tuning activity and membrane specificity of bacterial cytolysins"

**ONLINE SUPPLEMENTARY INFORMATION**

### SUPPLEMENTARY TABLES

**Supplementary Table 1.** Primer pairs used to generate CDC mutations. Mutated nucleotides are underlined and in bold.

| Mutation | Primer pairs (5’ to 3’) |
|---|---|
| **ILY** | |
| A221C | GCTGAAATCGGCGCCAAATTTGC**A**TTGCAGCTGGGACATACTTTG<br>CAAAGTATGTCCCAGCTGCAA**T**GCAAATTTGGCGCCGATTTCAGC |
| N404C | TTGGTTTCAATATAGTCCGTATTG**C**ACTGGATGGTTGCAATGCTATTATC<br>GATAATAGCATTGCAACCATCCAGT**G**CAATACGGACTATATTGAAACCAA |
| Y208C | CCGGCTCGCATGCAGT**G**CGAAAGCATCTCTGC<br>GCAGAGATGCTTTCG**C**ACTGCATGCGAGCCGG |
| D322C | GCAGGCGGCCATC**TG**TGCCGTCGTGAAAGGCGC<br>GCGCCTTTCACGACGGCA**CA**GATGGCCGCCTGC |
| L340C | GGAATACGAAAACATC**TGC**AAAAACACCAAAATCACG<br>CGTGATTTTGGTGTTTTT**GCA**GATGTTTTCGTATTCC |
| N342C | CGAAAACATCCTGAAA**TG**CACCAAAATCACGGC<br>GCCGTGATTTTGGTG**CA**TTTCAGGATGTTTTCG |
| E337A, N338A, K341A, N342A | GGTACGGAATACG**C**A**GC**CATCCTGG**GC**A**GC**CACCAAAATCACGG<br>CCGTGATTTTGGTG**GC**T**GC**CAGGATG**GC**T**G**CGTATTCCGTACC |
| Y336K, L340K | GCTGGTACGGAA**A**A**G**GAAAACATC**AA**GAAAAACACCAAAATCACGG<br>CCGTGATTTTGGTGTTTTTC**TT**GATGTTTTC**C**T**T**TTCCGTACCAGC |
| Y336W | GAAAGCTGGTACGGAAT**GG**GAAAACATCCTG<br>CAGGATGTTTTC**CC**ATTCCGTACCAGCTTTC |
| N338W | GGTACGGAATACGAA**TGG**ATCCTGAAAAACACC<br>GGTGTTTTTCAGGAT**CCA**TTCGTATTCCGTACC |
| L340W | CGGAATACGAAAACATC**TG**GAAAAACACCAAAATC<br>GATTTTGGTGTTTTTC**CA**GATGTTTTCGTATTCCG |
| N342W | CGAAAACATCCTGAAA**TGG**ACCAAAATCACGGC<br>GCCGTGATTTTGGT**CCA**TTTCAGGATGTTTTCG |
| **PLY** | |
| W278K, L282K | CCACAGACCGAG**GC**GAAGCAGATT**GC**GGACAATACGGAAG<br>CTTCCGTATTGTCC**GC**AATCTGCTTC**GC**CTCGGTCTGTGG |

| | |
|---|---|
| K279A, Q280A, D283A, N284A | CAGACCGAGTGG**GC**G**GC**GATTTTG**GC**C**GC**TACGGAAGTGAAGG<br>CCTTCACTTCCGTA**GC**G**GC**CAAAATC**GC**C**GC**CCACTCGGTCTG |

**Supplementary Table 2.** All CDC variants used in this study

| Variant | Mutations | Disulfide bonds | Source |
|---|---|---|---|
| **ILY** | | | |
| WT$^{charge+}$ | Y336K, L340K | | This study |
| WT$^{no\ charge}$ | E337A, N338A, K341A, N342A | | This study |
| WT$^{Y336W}$ | Y336W | | This study |
| WT$^{N338W}$ | N338W | | This study |
| WT$^{L340W}$ | L340W | | This study |
| WT$^{N342W}$ | N342W | | This study |
| prepore$^{\#}$ | I104C, G244C | C104-C244 | Boyd *et al*.[1] |
| prepore$^{L340C}$ | I104C, G244C, L340C | C104-C244 | This study |
| prepore$^{N342C}$ | I104C, G244C, N342C | C104-C244 | This study |
| prepore$^{charge+}$ | I104C, G244C, Y336K, L340K | C104-C244 | This study |
| HB1lock | A221C, N404C | C221-C404 | This study |
| HB1lock$^{charge+}$ | A221C, N404C, Y336K, L340K | C221-C404 | This study |
| HB1lock$^{no\ charge}$ | A221C, N404C, E337A, N338A, K341A, N342A | C221-C404 | This study |
| HB1lock$^{Y336W}$ | A221C, N404C, Y336W | C221-C404 | This study |
| HB1lock$^{N338W}$ | A221C, N404C, N338W | C221-C404 | This study |
| HB1lock$^{L340W}$ | A221C, N404C, L340W | C221-C404 | This study |
| HB1lock$^{N342W}$ | A221C, N404C, N342W | C221-C404 | This study |
| HB2lock | Y208C, D322C | C208-C322 | This study |
| HB2lock$^{charge+}$ | Y208C, D322C, Y336K, L340K | C208-C322 | This study |

| | | | |
|---|---|---|---|
| HB2lock$^{\text{no charge}}$ | Y208C, D322C, E337A, N338A, K341A, N342A | C208-C322 | This study |
| HB2lock$^{\text{Y336W}}$ | Y208C, D322C, Y336W | C208-C322 | This study |
| HB2lock$^{\text{N338W}}$ | Y208C, D322C, N338W | C208-C322 | This study |
| HB2lock$^{\text{L340W}}$ | Y208C, D322C, L340W | C208-C322 | This study |
| HB2lock$^{\text{N342W}}$ | Y208C, D322C, N342W | C208-C322 | This study |
| **PLY** | | | |
| WT$^{\text{charge+}}$ | W278K, L282K | | This study |
| WT$^{\text{no charge}}$ | K279A, Q280A, D283A, N284A | | This study |

#prepore refers to the ILY early prepore solved by our cryoEM reconstruction

**Supplementary Table 3.** Collection specifications of cryoEM data sets

| | **Data set 1** | **Data set 2** | **Data set 3** |
|---|---|---|---|
| Microscope | Titan Krios | Titan Krios | Titan Krios |
| keV | 300 | 300 | 300 |
| Camera | Falcon III | Falcon III | Falcon III |
| Collection mode | Linear | Linear | Linear |
| Pixel size (Å) | 1.4 | 1.4 | 1.4 |
| Stage tilt during collection | 0° | 0° | 30° |
| Number of frames | 39 | 39 | 39 |
| Integration time (s) | 1.00 | 1.00 | 1.22 |
| Total dose ($e^-/Å^2$) | 66.8 | 69.4 | 60.1 |
| Defocus range (µm) | -1.9 to -3.1 | -1.9 to -3.1 | -1.9 to -3.1 |
| Number of micrographs | 5202 | 5959 | 6878 |

**Supplementary Table 4.** Validation statistics for early prepore ILY-CD59 models

| | Central monomer | Oligomer |
|---|---|---|
| Molprobity score | 2.00 | 2.14 |
| Clash score | 10.13 | 14.71 |
| **Ramachandran plot** | | |
| Favourable | 92.44 | 92.44 |
| Allowed | 6.99 | 6.99 |
| Outliers | 0.57 | 0.57 |
| **Model correlation coefficient** | | |
| Global | 0.30 | 0.49 |
| Local* | 0.79 | 0.80 |

* Local model correlation coefficient determined by boxing density around model.

### SUPPLEMENTARY FIGURES

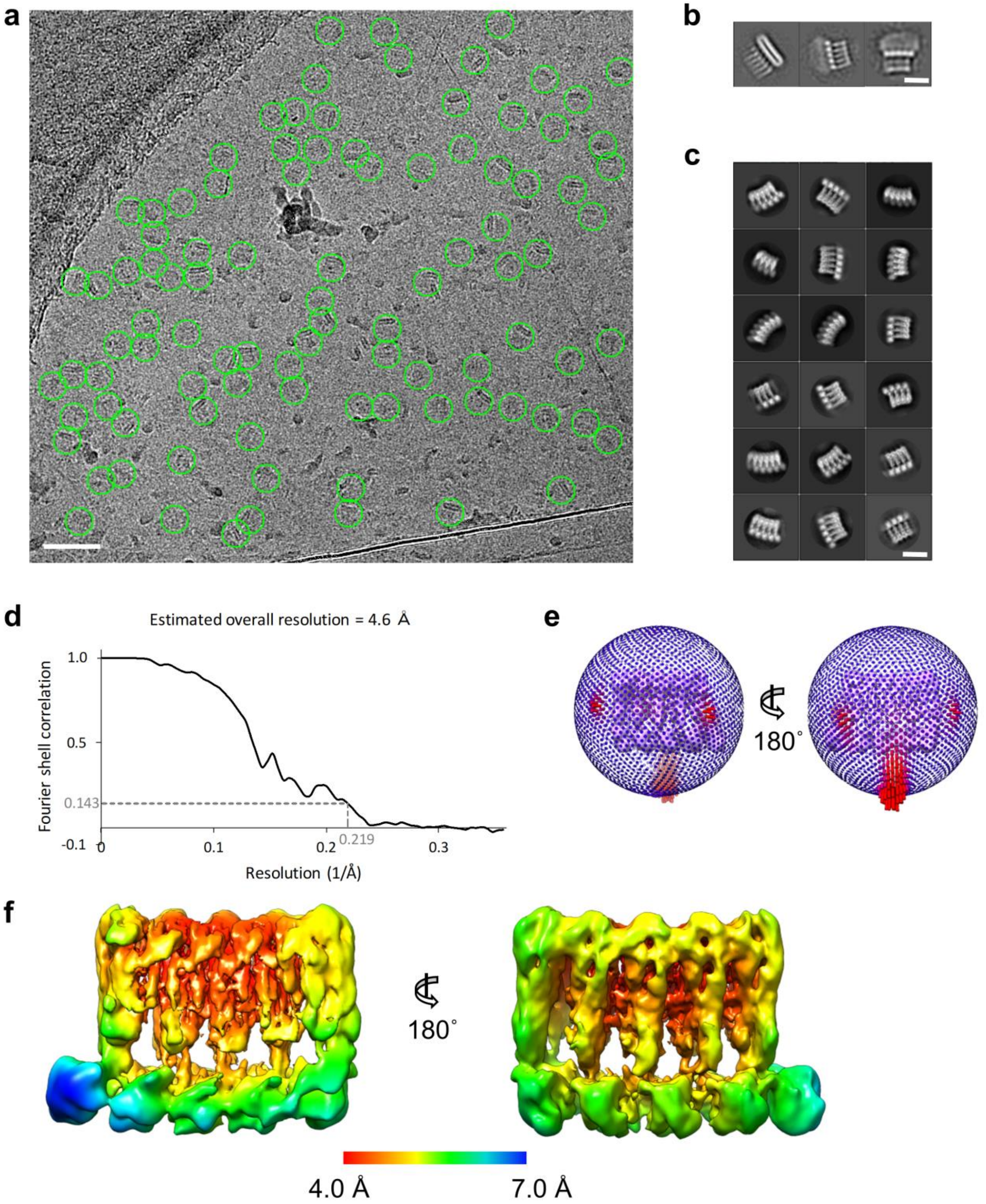


**Supplementary Figure 1.** CryoEM reconstruction of ILY early prepore in complex with CD59. (a) Representative electron micrograph of early prepore ILY on graphene-coated holey carbon grids. Particles circled in green. Scale bar, 500 Å. (b) Selected early 2D class averages that include nanodiscs, before particles were re-centered and masked to exclude nanodisc signal. Scale bar, 100 Å. (c) Selected 2D class averages from final classification. Scale bar, 100 Å. (d) Mask-corrected Fourier shell correlation (FSC) curve computed from unfiltered half-maps in RELION. (e) Angular distribution for the reconstruction. Height of the cylinder at each projection direction is proportional to the number of particle images, ranging from blue (fewer images) to red (more images). (f) Local resolution filtered map colored according to resolution, ranging from 4.0 to 7.0 Å. Data shown in (d-f) correspond to the final reconstruction obtained following the bottom right branch in Supplementary Figure 3 (purple).

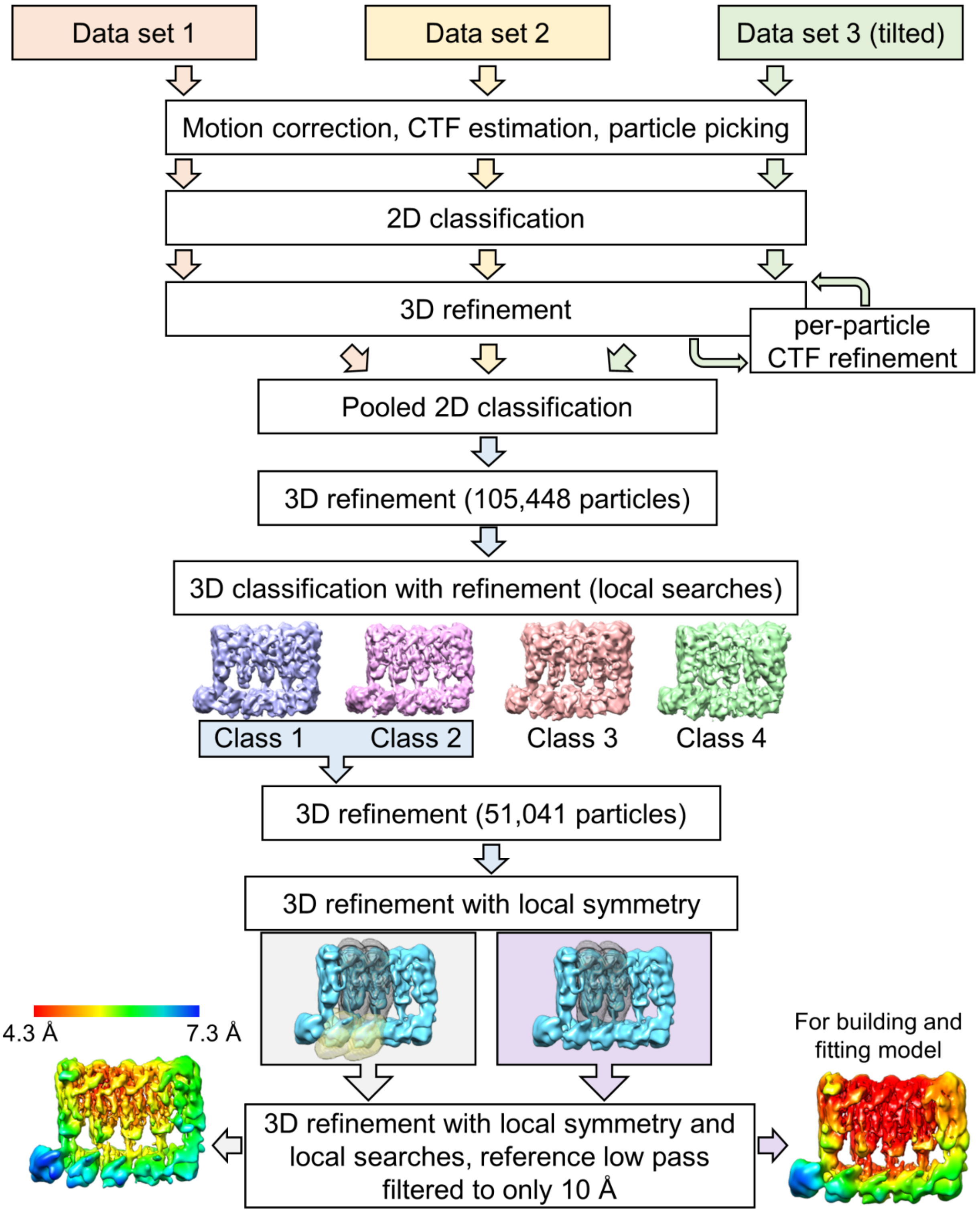


**Supplementary Figure 2.** Data analysis and reconstruction strategy for early prepore ILY oligomer. Particles were picked from three data sets (red, yellow, and green) whose micrograph movies were individually preprocessed (motion correction and CTF estimation). 2D classification of extracted particles from each data set removed poor particles, and the remaining particle sets were refined using an initial model generated in RELION. For Data set 3, per-particle CTF-refinement was performed to account for variation in z-height across the tilted micrographs, followed by another 3D refinement. All particles were then pooled (105,448 particles) and subjected to refinement to generate a consensus map. 3D classification with refinement separated these particles into 4 classes. Particles from Class 1 and 2 were combined (51,041 particles) and refined using local symmetry operators. In the right branch (purple background), the top regions of two monomers (grey) were assigned one set of symmetry operators. In the left branch, the bottom region of two monomers (yellow) were assigned an additional set of symmetry operators. A second round of 3D refinement with these respective symmetry operators was performed using local searches and a reference low pass filtered to 10 Å. The local resolution-filtered density reconstructions (bottom left and right) are shown colored according to the same resolution scale.

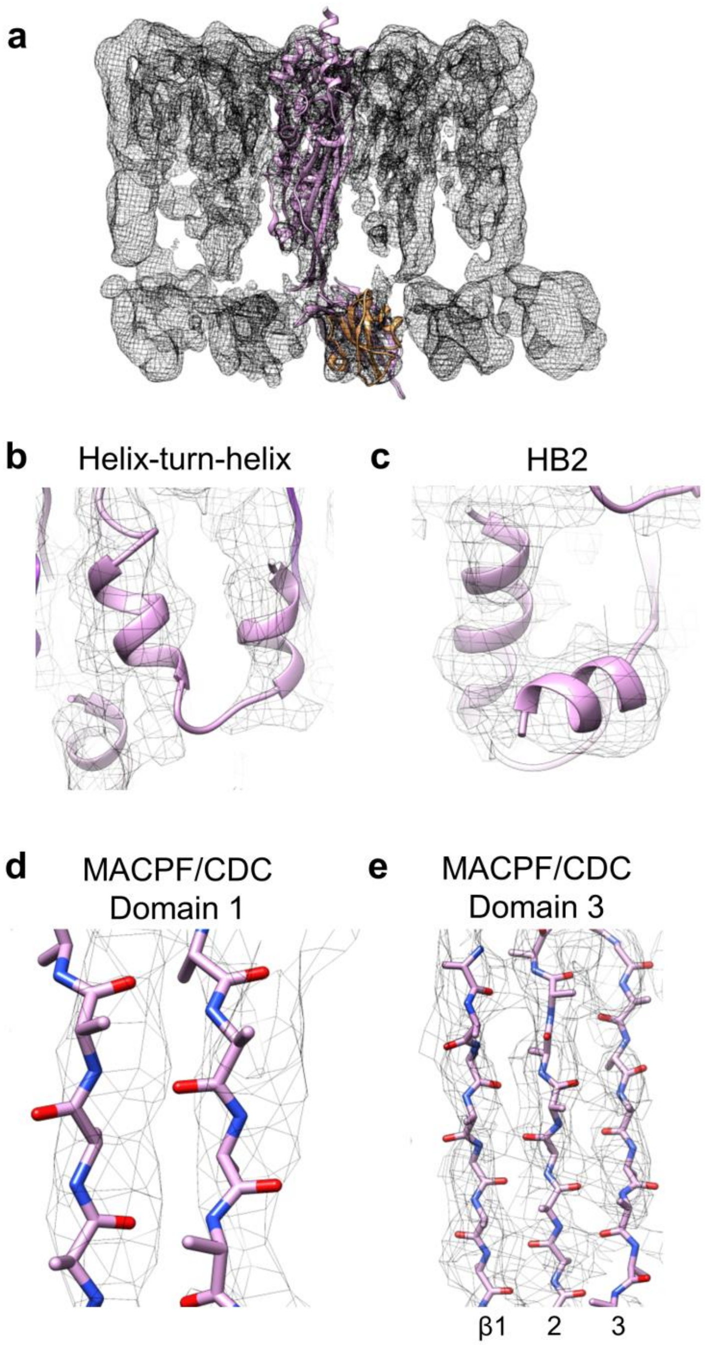


**Supplementary Figure 3.** Structural models of ILY (pink) and CD59 overlaid with the local resolution filtered cryoEM map (grey mesh). (a) Overlay of the ILY-CD59 early prepore model fit in the central monomer. (b) Close-up of the ILY helix-turn-helix motif (residues 356 -373) showing agreement of the map and model. (c) Close-up showing the quality of the fit for vertical and horizontal helices (h-helix) of HB2 (residues 315 to 342). (d) Quality of the map showing separation of β-strands within MACPF/CDC domain 1 (residues 258-261, 297-300). (e) Map and model overlay showing β-strands of MACPF/CDC domain 3 (residues 205-211, 249-256, and 301-308).

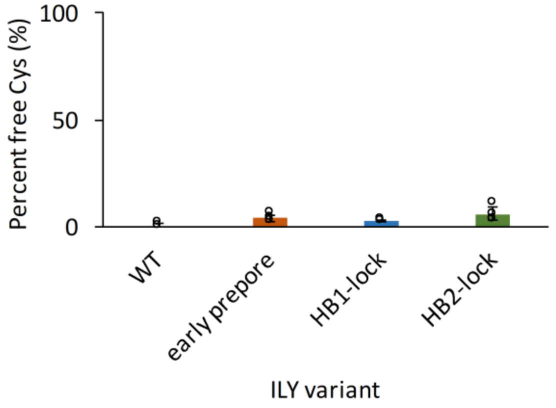


**Supplementary Figure 4.** Fluorescence-based cysteine accessibility assay. For wild type ILY (WT) and each disulfide-locked variant (early prepore, HB1-lock and HB2-lock), cysteine accessibility is expressed as a percentage of free cysteine (free cysteine/total cysteine), as determined by a standard curve. Individual data points shown as circles, n = 3. Error bars represent standard deviation.

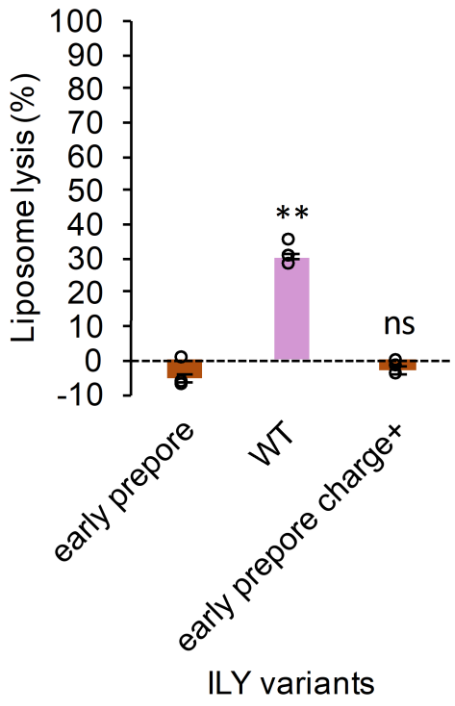


**Supplementary Figure 5.** Calcein-based liposome lysis assay. Wild type ILY (WT) and ILY early prepore variants were tested for their ability to lyse cholesterol containing liposomes decorated with CD59. Early prepore charge+ refers to the early prepore variant with h-helix mutations: Y336K, L340K. All experimental readings were normalized to a control reading of liposomes in buffer. Statistical significance displayed above each bar. Individual data points shown as circles, n = 3. Error bars represent standard deviation. P-value significance determined by one-way ANOVA with a Bonferroni post-test: ns, not significant; **, $p < 0.01$.
